# Microwell-enhanced optical rapid antibiotic susceptibility testing of single bacteria

**DOI:** 10.1101/2023.02.20.529233

**Authors:** Ireneusz Rosłoń, Aleksandre Japaridze, Stef Rodenhuis, Lieke Hamoen, Murali Ghatkesar, Peter Steeneken, Cees Dekker, Farbod Alijani

## Abstract

Bacteria that are resistant to antibiotics present an increasing burden on healthcare. To address this emerging crisis, novel rapid Antibiotic Susceptibility Testing (AST) methods are eagerly needed. Here, we present an optical AST technique that can determine the bacterial viability within one hour down to a resolution of single bacteria. The method is based on measuring intensity fluctuations of a reflected laser focused on a bacterium in reflective microwells. Using numerical simulations, we show that both refraction and absorption of light by the bacterium contribute to the observed signal. By administering antibiotics that kill the bacteria, we show that the variance of the detected fluctuations vanishes within one hour, indicating the potential of this technique for rapid sensing of bacterial antibiotic susceptibility. We envisage the use of this method for massively parallelizable AST tests and fast detection of drug resistant pathogens.

## Introduction

Antibiotic resistance is a major challenge for mankind^1^. Most common Antibiotic Susceptibility Testing (AST) methods are based on the detection of the growth rate of the pathogenic organisms or changes in the concentration of a marker molecule in solution^2^. In a clinical setting, the commonly adapted AST methods include measurement of the turbidity of the growth solution, its carbon dioxide content, or the diameter of the inhibition zone around an antibiotic disk^3^. These methods are the ’gold standards’ in the clinic, and have been used for over half of a century for their reliable and reproducible determination of Minimum Inhibitory Concentrations (MICs) of an antibiotic. However, these conventional methods are slow, due to their dependency on the growth rate of the pathogenic microorganism. As a result, AST methods typically require between 16 to 48 hours before any results can be obtained^4^, and for slowly growing pathogens, this waiting time may take up to weeks^5^. The slow growth also causes delays in determining the pathogen identity, typically done by MALDI-TOF mass spectrometry^6^ and therefore in prescribing the right antibiotic to patients. Numerous studies are being conducted to shorten the time between isolation of the pathogen and performing a rapid AST^7,8^. The target is to prescribe antibiotics on the same day as the diagnosis, which essentially means that AST should be performed within 8 hours or less^9^, and ideally even within an hour.

Various new methods have shown potential to obtain AST results faster than traditional methods^10,11,12^. These include full genome screening, microfluidic-based assays^8,13^, optical methods^14^, and nanomotion-based techniques^15,16^. Many of these emerging technologies obtained positive results in a laboratory setting, but face issues in clinical practice, where low cost and high throughput are key factors. Among them, optical phenotypic monitoring, which involves the detection of the motion of (groups of) motile pathogens, is interesting for its potential of performing AST within a few hours via tracing the changes in laser intensity when bacteria swim through a laser beam^14^. Although promising, the dependency of this technique to bacterial concentrations is a problem, since for low bacterial concentrations, long measurement times are often required to obtain sufficient statistics for determining the susceptibility of a microorganism to an antibiotic. Thus, methods are needed that enhance the likelihood of bacteria crossing the laser light.

Here, we present a new reflectometric read-out technique that performs AST on weakly trapped motile bacteria which greatly enhances the sensitivity of measurements, even to the level of detection of single bacteria. The technique detects bacterial motility via intensity variations in the laser light when a bacterium crosses the path of the laser beam that is reflected from the silicon surface. Recently we used suspended graphene drums to detect the nanomechancial motion of single bacteria adhered to its surface^16^. Here however, we show a different principle where we show that for motile bacteria we do not necessarily need a mechanical lever and bacteria viability can be detected solely by optical means. By patterning the surface with microwells that physically localize the bacteria within the laser focus, we increase the crossing event frequency. We demonstrate that the signal is significantly enhanced when cells are measured in proximity of a reflective surface and show that our reflectometric read-out system can be used for fast detection of the susceptibility of motile bacteria to antibiotics, opening a new route for rapid AST.

## Results

### Optical detection of single motile bacteria

The experiments were performed on silicon samples that were placed inside a cuvette containing motile MG1655 *E*.*coli* in LB medium. We recorded the intensity of the reflected 633 nm He-Ne red laser light (see Methods), that was focused on the silicon surface to a spot of 4 *μ*m in diameter, using a laser reflectometry setup as depicted in figure 1. The crossing of a bacterium through the focal region could be determined from the modulation of the intensity of the light that returned to the photodiode. Individual traces are normalized relative to their mean intensity, *V*_norm_(*t*) = (*V* (*t*)*/ < V* (*t*) *>*) − 1, where the brackets stand for the time average. The reflected laser intensity *V* (*t*) was measured for 30’, and the signal variance *σ*^2^ =*<* (*V*_norm_(*t*))^2^ *>* was used as a metric to compare various traces.

**Figure 1.**
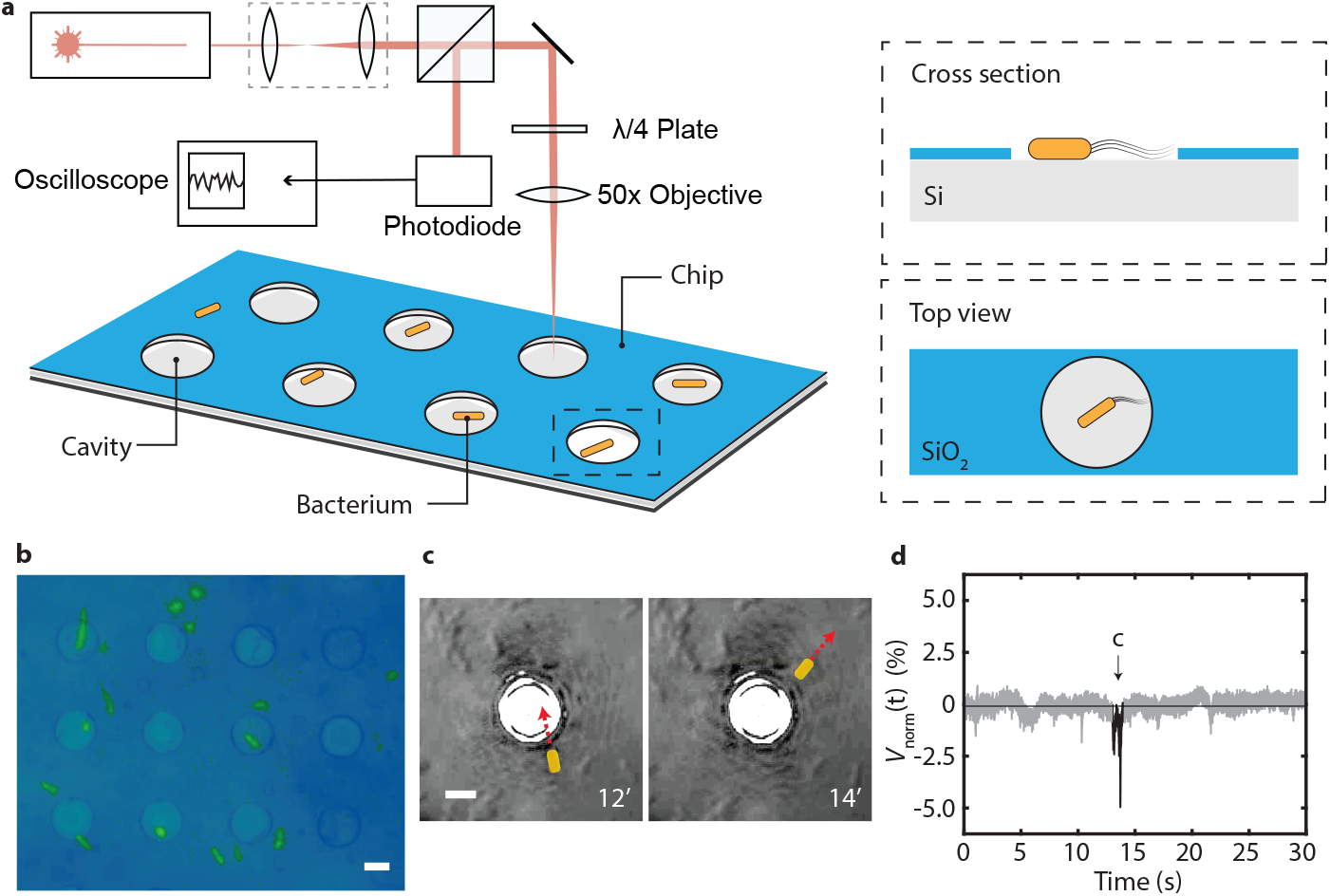
: Laser detection method for motile bacteria. a) Schematic illustration of interferometric read-out system used to localize bacteria in pre-patterned Si/SiO_2_ microwells with 8 *μ*m diameter and 285 nm depth. Bacteria on the patterned surface experience trapping, and stay longer inside the cavities than outside. b) Optical image of the fabricated microwells with fluorescent labelled bacteria, scale bar 5 micron. c) In-situ false-colored optical microscope image of microwells with a *E*.*coli* cell in LB suspension crossing laser focused on microwell. *E*.*coli* swimming path through microwell is indicated by a dotted line. Scale bar 5 micron. d) Drop in the detected signal during the bacterium crossing the laser path depicted on panel c (signal highlighted in black). Measurement performed at *E*.*coli* concentration OD=0.05.

When an *E*.*coli* bacterium passed through the laser focus, a sudden decrease in *V*_norm_(*t*) was recorded. Figure 1c-d as well as Supplementary Video 1 present an example, where we simultaneously acquired the signal intensity and performed optical tracking of a cell during such an event. We observed fluctuations in the read-out signal when a bacterium crossed the laser focus from the bottom to the upper right. Such fluctuations were not observed when bacteria were absent (see Supplementary Note 1). In the presence of motile bacteria, the fluctuations amounted up to a 10 % of the total light intensity incident on the photodiode (figure S1).

To enhance the frequency of events, we introduced micro patterned silicon substrates, where we used microwells (see fig. 1a) to localize the bacteria. Such microwells are known to be able to trap the bacteria for prolonged time^17^, and can thus be used to maintain them in close vicinity of both the laser focus and the silicon surface without impeding their motility. While focusing the laser onto a microwell, we observed that the signal appeared in prolonged ’bursts’: periods of increased fluctuations that were followed by a period of relative rest. As a result, individual traces in this case showed extended fluctuations (see Fig. 2c as an example). We performed the three different experiments in LB growth medium: without bacteria on bare silicon (as a control experiment), with bacteria on bare silicon, and with bacteria on micro patterned cavities. Typical traces at *E*.*coli* concentration with optical density OD=0.2 are shown in Figs. 2a-c. A statistical analysis of the variance of such traces is shown in figure 2d.

**Figure 2.**
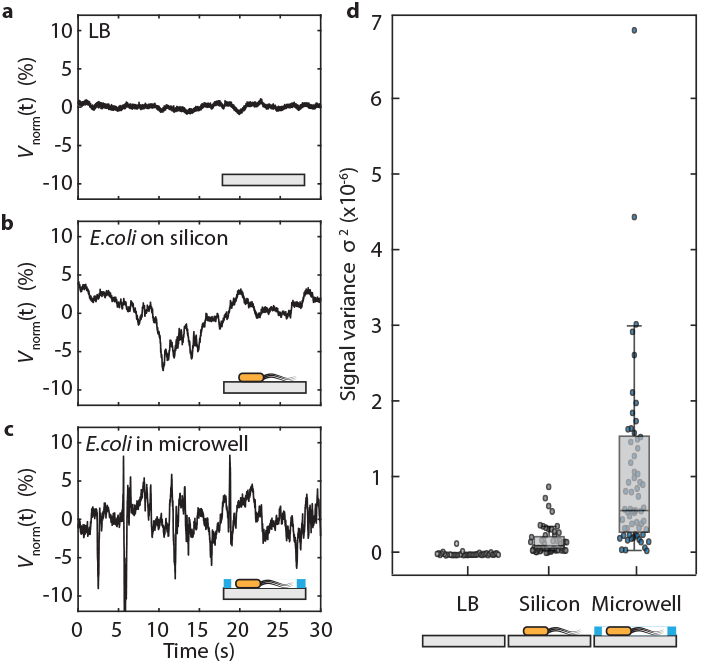
: Experiments on silicon surface and in microwells. a-c) Typical signals recorded in three cases: a) a control measurement on flat silicon with Lysogeny Broth (LB) without bacteria; b) on flat silicon surface with LB containing bacteria, and c) on a micro-well containing bacteria in LB. d) Signal variance for these three cases: on reflective bare silicon with LB (*n* = 43), same in the presence of bacteria (*n* = 54), and on bacteria in micro patterned wells (*n* = 67). Signal fluctuations appear only when bacteria are present, and signals are prolonged when the surface is patterned with microwells.

When we performed the experiment on flat silicon, we noted that bacteria crossed the laser focus only rarely during the 30’ observation windows, and traces for bacteria on bare silicon accordingly showed a variance that is only slightly higher than that in the control experiment. Indeed, in Figure Fig. 2b similar to Figs. 1c-d we saw a single crossing event where only a spike in the intensity signal could be observed. The measurements on microwell substrates however, showed on average a significantly higher variance. The probability for detecting laser intensity variations due to bacteria crossing the laser path was thus significantly enhanced by performing the experiment in the microwells where the bacteria got trapped. The localization of the bacteria inside the microwells was further confirmed by microscopic imaging on transparent PDMS samples with the same geometry as the silicon (see figure S2), in which we observed a 50% higher probability of imaging a bacterium inside a well than outside of it.

### Dependence of read-out signal on bacterium size

The finding that signal fluctuations are due to bacteria crossing the laser path, suggests that the strength of the read-out signal could be dependent on the size of the bacteria. Since the laser beam is larger than the bacterium diameter, changing the bacterium shape or size can be expected to cause a different light refraction and absorption by the cell. In order to test this hypothesis, we measured the signal of shape- and size-manipulated *E*.*coli* cells. We grew the bacteria in the presence of low doses of A22^18^ or Cephalexin^19^, which changed the bacterial cells into spherical and tubular shapes, respectively.

Figure 3a-b *E*.*coli* compares the data for bacteria with different sizes. Normal rod-shaped *E*.*coli* cells had a length of 3.4 ± 0.6 micron (*n* = 51) and a width of 1.0 ± 0.1 micron, while spherical A22 exposed cells had a diameter of 4.2 ± 0.6 micron (*n* = 34). The cells that were exposed to Cephalexin grew along the longitudinal axis, forming tubular shapes with a length of 10 ± 2 (*n* = 43) microns. As expected, changes in cell shape influenced the observed signal fluctuations, with larger cells generating larger signal fluctuations.

**Figure 3.**
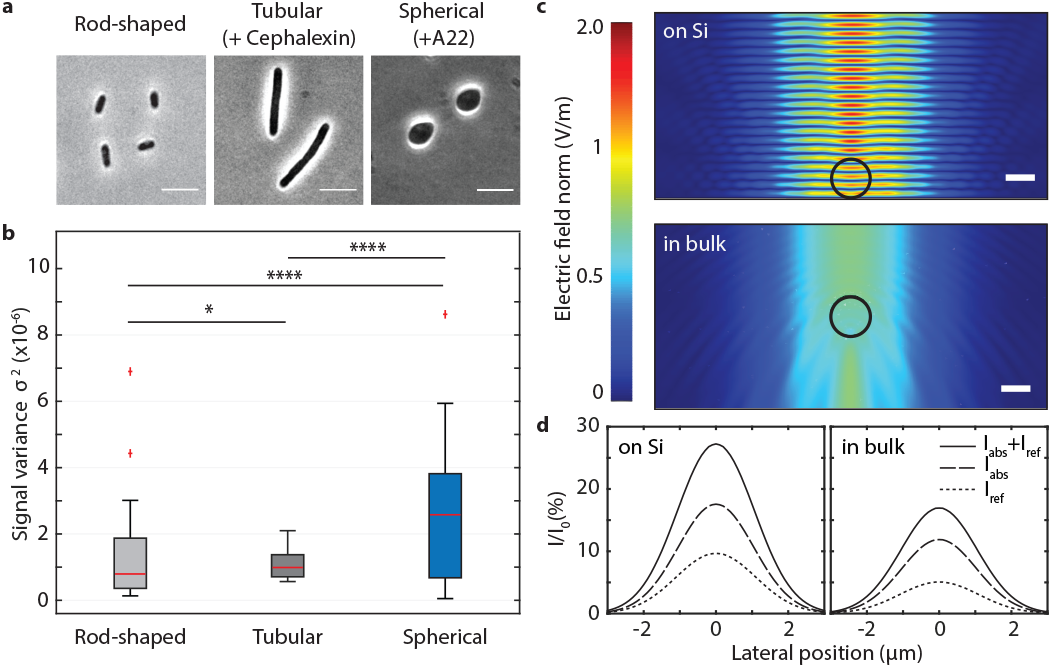
: Cell size and position determine the amount of light that is attenuated. a) Phase contrast images images of the three different sized *E*.*coli* cells used in our experiments. Scale bars are 5 *μ*m. b) Microwell measurements were performed on bacteria with different cell sizes: normal rod shaped cells (3.4 micron in length), long tubular cells (10 micron in length), and spherical cells (4.2 micron diameter). Boxplot whiskers extend to maximum 1.5 times the inter-quartile distance and outliers are indicated with crosses. Red horizontal line represents the median values and significance is expressed using the asterisk convention. c) Simulated electric field amplitude *E* of a focused 633 nm laser beam around an *E*.*coli* cell (indicated by a black circle) on a perfectly reflective silicon surface (top graph), and freely suspended in LB medium (bottom graph). The laser is incident from the top of the plot and the silicon surface is at the bottom of the top plot. Scale bar is 1 *μ*m and laser beam waist is 4*μ*m, similar to experimental conditions. d) Fraction of the incident laser power that is attenuated (*I*_abs_ + *I*_ref_) by absorption (*I*_abs_) and refraction (*I*_ref_) as a function of the lateral position of the cell, both in case of on the reflective silicon surface (left) and freely suspended in the growth medium (right).

### The role of position, absorption, and refraction in signal detection

Light intensity fluctuations can be attributed to two main sources: first, the absorption of the laser light by a bacterium, and second, refraction of light at the boundary of the bacterium, caused by the difference in the refractive indices of the cell and the surrounding medium. Typical values of the refractive index for *E*.*coli* are 1.39 ± 0.05, while values for LB medium have been reported as 1.335 ± 0.03^20,21,22^. Light travelling through a bacterium is absorbed more than in the surrounding liquid, a property which is typically used in cell counting experiments by optical density (OD) measurements^23,24,25^. For *E*.*coli* cells we used an attenuation coefficient of *μ* = 1.1 *×* 10^5^ m^−1^ (see Methods). Using these estimates, we performed COMSOL finite element simulations of Maxwell equations to explore the influence of a bacterium on the optical field and to find what portion of light is attenuated by a single bacterium passing through a focused laser beam (see also figure S3). In these simulations, the distortion and intensity change of a Gaussian beam with a waist diameter of 4 *μ*m was calculated in LB medium, both without and in the presence of an *E*.*coli* bacterium.

Figure 3c shows the electric field amplitude *E* for both the case where the bacterium is near the silicon surface and for the case that it is far from it. Simulations were performed in 2D and the cell is represented by a black circle. The interference between incident and reflected light waves results in a prominent standing wave near the silicon surface. In the absence of the silicon substrate there is no standing wave and the refraction caused by the bacterium can be observed clearly. These calculations were repeated for various positions of the cell relative to the laser focal position, to simulate a bacterium swimming through the center of the beam (see also Supplementary Note 3). Furthermore, we calculated the absorption of the electric field *I*_abs_ by the bacterium for all lateral positions by integrating the power loss *p* over the cell area *A, I*_abs_ =∯_*A*_ *p* d*A*, as shown in figure 3d as a percentage of the total incoming optical power *I*_0_. We also computed the amount of light *I*_ref_ that is refracted by an angle greater than 45 degrees (the limit due to the numerical aperture of the lens), i.e. the light not returning to the detector again as a function of the lateral position of the bacterium. Far outside of the laser beam, obviously, the bacterium does not absorb nor refract light. In the center of the laser beam though, up to 18 % of the incoming light is absorbed and up to 10 % is refracted if the bacterium is on the silicon surface, which indicates that both absorption and refraction by the cells play a role. These numbers are similar to the value in experiments, where typical oscillations are 10% and the highest peak to-peak variations that we observed were 20% of the total signal amplitude. Notably, in most experiments, the cells did not cross the beam exactly in the center of the laser focus and hence the experimental values are lower than simulated.

The bacteria close to the silicon surface yielded a signal that is about twice higher than that of bacteria that were swimming freely in bulk LB (see figure 3d). Since the laser beam is focused with an 0.55 NA objective to a 4 *μ*m spot, creating of conical bundle with a 46^*◦*^ angle, we are mostly sensitive to bacteria close to the focal point. The light beam quickly spreads wider away from the surface. For example, at a height of 10 *μ*m away from the surface, the light beam cross section is already 12.7 *μ*m, i.e. about 10 times larger than the cross section of a typical bacterium, and the signal from a bacterium crossing far away from the focal point is reduced by ten fold. Accordingly, bacteria need to be close to the focal point to be detected. Guiding or trapping bacteria near the surface and laser focus can thus improve the read-out signal in addition to increasing the event frequency of bacteria passing the laser light.

### Antibiotic susceptibility of single cells

Finally, we explored if this method can be applied for testing the efficacy of antibiotics. We compared the signal of live bacteria on bare silicon and on patterned microwells, to the signal of the bacteria after exposure to various antibiotics. We tested chloramphenicol, an antibiotic that blocks protein synthesis^26^, and ciprofloxacin, an antibiotic that blocks the activity of DNA gyrases^27^. Importantly these antibiotics don’t affect the morphology, the size and shape, of the bacteria (figure S4).

Figure 4 shows the signal variances before and after administering the antibiotic for both cases. One hour after the addition of antibiotics, there was a significant drop in the signal for both antibiotics. For the data on a silicon surface, however, no significant change could be observed after addition of the chloramphenicol.

**Figure 4.**
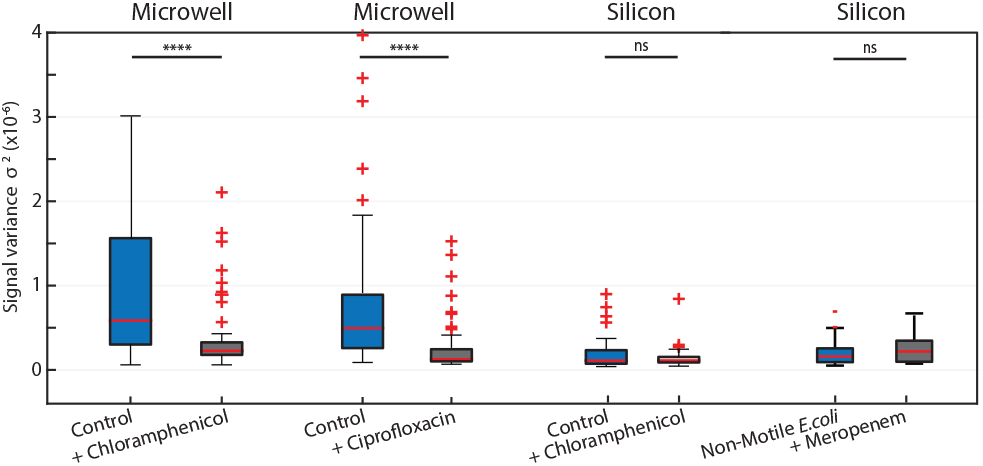
: Effect of antibiotics on the observed signal amplitude. Signals before and after administering various antibiotics. 1h after administering Chloramphenicol (34*μ*g/ml), 1.5h after administering Meropenem (50*μ*g/ml) and 3h after administering Ciprofloxacin (20*μ*g/ml). On the etched microwells (n = 67), after administering antibiotics, chloramphenicol (n=67) or ciprofloxacin (n=200), a significant drop of the initial signal (*p <* 10^*−*5^, ****) can be observed for susceptible bacteria. For measurements on bare silicon (n = 54) no significant difference (*p* = 0.94, ns) was measured after exposure to the antibiotic (n = 54). Also *E*.*coli motAB* non-motile cells were tested, and no statistical difference between signal from on non-motile ((*n* = 71) and antibiotic treated cells (*n* = 82) was observed. Boxplot whiskers extend to maximum 1.5 times the interquartile distance and outliers are indicated with crosses. Red horizontal line represents the median values. Measurements are compared using a two-tailed Wilcoxon ranksum test.

To test the efficacy of the technique in detecting antibiotic resistance, we also performed an additional experiment on *E. coli* with *KanR* resistance gene^28^. We exposed these resistant cells to kanamycin^29^, an antibiotic that inhibits protein synthesis, but we did, as expected, not observe a change in the variance of the signal after administering the antibiotic (see figure S5). Even after several hours of incubation the signal stayed unchanged, demonstrating that the technique is able to demonstrate not only susceptibility to antibiotics but also the resistance of bacteria against them.

In a recent study, wide field optical microscopy was employed for the detection of movement of single bacteria^30^. It reported observation of motion even for non-motile bacterial species such as *S. aureus*, which diminished after treatment with antibiotic. To verify this hypothesis, we tested E.coli motAB non-motile cells and did not observe a significant change in signal amplitude before and after antibiotic treatment (see Figure 4). In fact, we observed signals that were of the same magnitude as background (see figure S6). We note that we perform a Wilcoxon rank-sum test in order to compare the two non-normal distributions, whereas the recent study performed a student’s t-test. This might explain the difference in the conclusions of the works.

## Discussion

We presented an optical detection technique to measure the viability of single motile bacterial cells. Our method is based on the fluctuations of a laser signal when bacteria run through its focal plane. To extend the time during which a bacterium motility can be measured in the laser spot, we introduced microwells in the silicon surface with 285 nm depth and 8*μ*m diameter. Because the bacteria are trapped at these predetermined microwell spots, the bacteria stay longer near the laser spot (figure 1a), and more events can be observed during the measurement window, i.e. larger signals are collected. The cavity dimensions are chosen to be comparable with the size of bacteria. A relatively shallow cavity depth minimizes optical aberrations that would be introduced by the sidewalls. To yet further increase the throughput of this method, a more elaborate chip design can be conceived. One could for example guide bacteria towards the laser focus by channels or mazes^31^, which would allow samples at lower concentrations to be used for detection. Optimization of the trap depth might also aid measurements see for example figure S6 where the duration of trapping events is prolonged although too deep traps will impact readout quality adversely and might limit the natural motility of the bacteria.

Next to our experimental observations, we performed numerical studies, and concluded that the variations in the reflected signal can be explained by a combination of refraction and absorption of the laser light by the *E. coli* bacteria. Peak variations in signal during experiments (up to 20%) were of comparable magnitude as the maximum variations that were calculated from the simulations (maximum 28%). Our finite-element simulations showed that bacterial motion resulted in larger signal fluctuations near a reflective surface than in the free volume. The simulations provide a better understanding of the optimal conditions for optical detection.

The detected signal in measurements of the bacteria described here is directly linked to the motility of the pathogens, which vanishes upon exposure to antibiotics. We believe that the high-speed nature of our technique will be helpful for developing rapid diagnostic tools for detection for AST of motile pathogens. For example, in urinary tract infections by *E*.*coli* (which accounts to 75% of infections)^32^, we envisage our technique to be highly efficient. It is important to highlight that we could detect the susceptibility and resistance to antibiotics in less than an hour, which is significantly quicker than existing detection techniques based on growth rate of bacteria that typically take days^33^. We are confident that the current results provide a good base to further accelerate the development of next-generation AST tests.

## Limitations of the study

The technique described in this work is ineffective for non-motile pathogens. Therefore, in case of an infection with a non-motile pathogen, alternative methods must be used for AST.

## Acknowledgements

We acknowledge Michiel Otte and Le-Vaughn Naarden for their help in performing measurements with deep cavities. Financial support was provided from the European Union’s Horizon 2020 research and innovation programme under ERC starting grant ENIGMA (no. 802093, F.A. and I.E.R.), ERC PoC GRAPHFITI (no. 966720, F.A. and A.J.), Dutch Research Council (NWO) take-off grant, Graphene Flagship (grant nos. 785219 and 881603, P.S.) and the ERC Advanced Grant Looping-DNA (no. 883684, C.D.), as well as NWO/OCW as part of the NanoFront and BaSyC programmes and by the Swiss National Science Foundation (grant no. P300P2 177768, A.J.).

## Author contributions

I.E.R., A.J, F.A. conceived the idea. I.E.R., A.J. and S.R. collected the data and performed the interferometry experiments. I.E.R. constructed the setup and performed the simulations. A.J. and L.H performed the bacterial manipulation. All authors designed the experiments. The project was supervised by F.A, M.G, C.D., and P.G.S. All authors contributed to the data analysis, interpretation of the results and writing of the manuscript.

## Data availability

The data that support the findings of this study are available from the corresponding authors upon request.

## Methods

### Sample preparation

All experiments we performed on MG1655(+IS1) *E. coli* cells, described earlier^34^. Experiments with Kanamycin resistant *E*.*coli* cells were performed on MG1655(*kanR*) cells described earlier^16^. The *E. coli* cells, were grown in LB medium overnight at 30^*◦*^C to reach the late exponential phase. The next day before performing experiment, the culture was refreshed (1:100 volume) for 2.5 hours on fresh LB medium at 30^*◦*^C reach an optical density (OD600) OD=0.2. The chamber was filled with the solution at this concentration, unless stated otherwise, and left for 15 minutes horizontal position to deposit the bacteria on the surface. For experiments where antibiotics were used, antibiotics were dissolved in LB and incubated with bacteria for 1h. Chloramphenicol was used at 34*μ*mg/ml, Ciprofloxacin at 20*μ*g/ml, Kanamycin at 25*μ*g/ml and Meropenem at 50*μ*g/ml final concentration. An optical microscope (Keyence VHX-7000) was used to inspect the sample. The chamber was placed in the interferometric setup that was equipped with Attocube ECSx5050 nano positioners that allow automated scanning. The motion of the bacterium caused changes in the optical path, that were monitored by a photodiode and an oscilloscope (Rohde & Schwarz RTB2004). At each measured point on the substrate, a trace was recorded for 30 seconds with 50’000 data points. The measurements were performed in an air-conditioned room with a temperature of 21^*◦*^C. The substrates were either 5×5 mm^2^ silicon chips, or 5×5 mm^2^ silicon chips with a 285 nm layer of silicon oxide. The latter were patterned with circular cavities by a reactive ion etch, where silicon acted as a stop layer, creating cavities with a diameter of 8 *μ*m, described earlier^16^.

### Bacterial shape manipulation

In order to grow the *E*.*coli* cells into spherical shapes, low doses of the A22 drug were added to the to LB. On the day of the experiment, the cell culture was refreshed (1:100 volume) in the presence of A22 drug (5*μ*g/ml final concentration) for 1.5 hours on fresh LB medium at 30^*◦*^C reach an optical density (OD600) between OD=0.2–0.3. A22 inhibits the MreB polymerization, thereby disrupting the typical rod shape of *E. coli* ^18^. These spherical cells remain physiologically active and can replicate and divide^35,36^. In order to grow the cells into tubular shapes, low doses of cephalexin drug (25*μ*g/ml final) were added to the to LB and cells were grown for 1 hours on fresh LB medium at 30^*◦*^C. Cephalexin blocks cell division but allows cells to grow in length^19^.

### Optical Microscopy

To measure the sizes of *E*.*coli* cells we used Nikon Ti-E microscope with a 100X CFI Plan Apo Lambda Oil objective with an NA of 1.45 equipped with a phase ring. Images were captured by Andor Zyla USB3.0 CMOS Camera.

### Statistics

Since the data reported in the paper are not normally distributed, we relied on non-parametric tests for statistics. We represent the median and quartiles of data in boxplots, in accordance with the use of non-parametric tests. We use a rank sum test for comparison between measurement sets. We used MATLAB’s built-in functions for statistical analysis. All statistical tests were two-sided. On all figures, the following conventions are used: not significant (NS) 0.05 *<* P, *0.01 *<* P *<* 0.05, **0.001 *<* P *<* 0.01, ***0.0001 *<* P *<* 0.001, ****P *<* 0.0001. We report a significant difference in results if P *<* 0.01.

### Laser interferometry

A red laser (*λ*_red_ = 632.8 nm) focused with a 4 *μ*m spot size on the sample was used for detection of the amplitude of the cell motion, where the position-dependent optical absorption of the cell results in an intensity modulation of the reflected red laser light, that was detected by a photodiode^37^. The incident red laser power was 3 mW.

### Calculation of linear attenuation coefficient

The optical density (OD) of a sample is defined as the logarithm of the ratio between the incident and transmitted laser power, that is: *OD* = *log*_10_(*I*_1_*/I*_0_). This means that at *OD* = 1, a fraction *x* = 0.1 of the incident light is transmitted. A measurement of *OD* = 1 corresponds to approximately 10^9^ bacteria /mL in a 1 cm cuvette^38^.The fraction of light *x* that is transmitted by a single bacterium can thus be expressed as 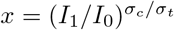, where *σ*_*c*_ is the physical cross section of the cuvette and *σ*_*t*_ is the total cross section of *n* bacterial cells in suspension with each a physical cross section *σ*, i.e. *σ*_*t*_ = *nσ*. We wish to compute the linear attenuation coefficient *μ*, which relates the transmitted laser power to the distance *d* travelled through a bacterium by the following expression: *I*(*d*) = *I*_0_ *e*^*−μd*^. This can be rewritten into *μ* = − *ln*(*x*)*/d*. From the measured physical cross section of a single bacterium (*A ≈* 1 × 2*μ*m^2^)^39^ and the cross section of the cuvette (*A* = 1 cm^2^), we find that a single bacterium absorbs around *x* = 11% of the incoming light and an attenuation coefficient of *μ* =− *ln*(*x*)*/d* = 1.1 × 10^5^ m^*−*1^ for *E*.*coli* cells with average diameter *d* = 1*μ*m.

### Data processing

The signal obtained from the photodiode voltage due to the variations in reflected intensity of the red laser is recorded by an oscilloscope. The time trace of the photodiode voltage *V*_pd_(t) was normalized by division over its average, *V*_norm_(*t*) = *V*_pd_(*t*)*/ < V*_pd_(*t*) *>*, after which a linear fit was subtracted from the data to eliminate the effects of drift during the measurements.

## References

1. J. O’Neill (2016). Tackling drug-resistant infections globally: final report and recommendations.

2. L. B. Reller, M. Weinstein, J. H. Jorgensen, and M. J. Ferraro (2009). Antimicrobial Susceptibility Testing: A Review of General Principles and Contemporary Practices. Clinical Infectious Diseases 49.11, pp. 1749–1755. issn: 1058-4838. doi: 10.1086/647952.

3. H. Leonard, R. Colodner, S. Halachmi, and E. Segal (2018). Recent advances in the race to design a rapid diagnostic test for antimicrobial resistance. ACS sensors 3.11, pp. 2202–2217.

4. S. Puttaswamy, S. K. Gupta, H. Regunath, L. P. Smith, and S. Sengupta (2018). A comprehensive review of the present and future antibiotic susceptibility testing (AST) systems. Arch Clin Microbiol 9.3.

5. L. Hall, K. P. Jude, S. L. Clark, and N. L. Wengenack (2011). Antimicrobial susceptibility testing of Mycobacterium tuberculosis complex for first and second line drugs by broth dilution in a microtiter plate format. JoVE (Journal of Visualized Experiments) 52, e3094.

6. T.-Y. Hou, C. Chiang-Ni, and S.-H. Teng (2019). Current status of MALDI-TOF mass spectrometry in clinical microbiology. journal of food and drug analysis 27.2, pp. 404–414.

7. G. D. Kaprou, I. Bergšpica, E. A. Alexa, A. Alvarez-Ordóñez, and M. Prieto (2021). Rapid methods for antimicrobial resistance diagnostics. Antibiotics 10.2, p. 209.

8. Ö. Baltekin, A. Boucharin, E. Tano, D. I. Andersson, and J. Elf (2017). Antibiotic susceptibility testing in less than 30 min using direct single-cell imaging. Proceedings of the National Academy of Sciences 114.34, pp. 9170–9175.

9. A. van Belkum, T. T. Bachmann, G. Lüdke, J. G. Lisby, G. Kahlmeter, A. Mohess, K. Becker, J. P. Hays, N. Woodford, K. Mitsakakis, et al. (2019). Developmental roadmap for antimicrobial susceptibility testing systems. Nature Reviews Microbiology 17.1, pp. 51–62.

10. A. van Belkum, C.-A. D. Burnham, J. W. Rossen, F. Mallard, O. Rochas, and W. M. Dunne (2020). Innovative and rapid antimicrobial susceptibility testing systems. Nature Reviews Microbiology 18.5, pp. 299–311.

11. F. Zhang, J. Jiang, M. McBride, X. Zhou, Y. Yang, M. Mo, J. Peterman, T. Grys, S. E. Haydel, N. Tao, et al. (2021). Rapid Antimicrobial Susceptibility Testing on Clinical Urine Samples by Video-Based Object Scattering Intensity Detection. Analytical Chemistry 93.18, pp. 7011–7021.

12. Z. A. Khan, M. F. Siddiqui, and S. Park (2019). Current and emerging methods of antibiotic susceptibility testing. Diagnostics 9.2, p. 49.

13. R. M. Humphries (2020). Update on susceptibility testing: genotypic and phenotypic methods. Clinics in Laboratory Medicine 40.4, pp. 433–446.

14. I. Bennett, A. L. Pyne, and R. A. McKendry (2020). Cantilever sensors for rapid optical antimicrobial sensitivity testing. ACS sensors 5.10, pp. 3133– 3139.

15. L. Venturelli, A.-C. Kohler, P. Stupar, M. I. Villalba, A. Kalauzi, K. Radotic, M. Bertacchi, S. Dinarelli, M. Girasole, M. Pešić, et al. (2020). A perspective view on the nanomotion detection of living organisms and its features. Journal of Molecular Recognition 33.12, e2849.

16. I. E. Rosloń, A. Japaridze, P. G. Steeneken, C. Dekker, and F. Alijani (2022). Probing nanomotion of single bacteria with graphene drums. Nature Nanotechnology, pp. 1–6.

17. C. Probst, A. Grünberger, W. Wiechert, and D. Kohlheyer (2013). Polydimethylsiloxane (PDMS) Sub-Micron Traps for Single-Cell Analysis of Bacteria. Micromachines 4.4, pp. 357–369. ISSN: 2072-666X. doi: 10.3390/mi4040357.

18. A. Varma and K. D. Young (2009). In Escherichia coli, MreB and FtsZ direct the synthesis of lateral cell wall via independent pathways that require PBP 2. Journal of bacteriology 191.11, pp. 3526– 3533.

19. N. Maki, J. E. Gestwicki, E. M. Lake, L. L. Kiessling, and J. Adler (2000). Motility and chemotaxis of filamentous cells of Escherichia coli. Journal of Bacteriology 182.15, pp. 4337–4342.

20. B. Gul, S. Ashraf, S. Khan, H. Nisar, and I. Ahmad (2021). Cell refractive index: Models, insights, applications and future perspectives. Photodiagnosis and Photodynamic Therapy 33, p. 102096.

21. K. Stevenson, A. F. McVey, I. B. Clark, P. S. Swain, and T. Pilizota (2016). General calibration of microbial growth in microplate readers. Scientific reports 6.1, pp. 1–7.

22. A. E. Balaev, K. Dvoretski, and V. A. Doubrovski (2002). Refractive index of Escherichia coli cells. In Saratov Fall Meeting 2001: Optical Technologies in Biophysics and Medicine III. Vol. 4707. SPIE, pp. 253–260.

23. P. Liu, L. Chin, W. Ser, T. Ayi, P. Yap, T. Bourouina, and Y. Leprince-Wang (2014). An optofluidic imaging system to measure the biophysical signature of single waterborne bacteria. Lab on a Chip 14.21, pp. 4237–4243.

24. P. Y. Liu, L. Chin, W. Ser, H. Chen, C.-M. Hsieh, C.-H. Lee, K.-B. Sung, T. Ayi, P. Yap, B. Liedberg, et al. (2016). Cell refractive index for cell biology and disease diagnosis: past, present and future. Lab on a Chip 16.4, pp. 634–644.

25. Y. P. Liu (2016). “Refractive Index Distribution of Single Cell and Bacterium Usingan Optical Diffraction Tomography System”. PhD thesis. Université Paris-Est.

26. O. Pongs, R. Bald, and V. A. Erdmann (1973). Identification of chloramphenicol-binding protein in Escherichia coli ribosomes by affinity labeling. Proceedings of the National Academy of Sciences 70.8, pp. 2229–2233.

27. D. C. Hooper and G. A. Jacoby (2016). Topoisomerase inhibitors: fluoroquinolone mechanisms of action and resistance. Cold Spring Harbor perspectives in medicine 6.9, a025320.

28. E. Bremer, T. Silhavy, and G. Weinstock (1985). Transposable lambda placMu bacteriophages for creating lacZ operon fusions and kanamycin resistance insertions in Escherichia coli. Journal of bacteriology 162.3, pp. 1092–1099.

29. H. Yamaki and N. Tanaka (1963). Effects of protein synthesis inhibitors on the lethal action of kanamycin and streptomycin. The Journal of Antibiotics, Series A 16.6, pp. 222–226.

30. M. I. Villalba, E. Rossetti, A. Bonvallat, C. Yvanoff, V. Radonicic, R. G. Willaert, and S. Kasas (2023). Simple optical nanomotion method for single-bacterium viability and antibiotic response testing. Proceedings of the National Academy of Sciences 120.18, e2221284120.

31. P. Galajda, J. Keymer, P. Chaikin, and R. Austin (2007). A wall of funnels concentrates swimming bacteria. Journal of bacteriology 189.23, pp. 8704– 8707.

32. A. L. Flores-Mireles, J. N. Walker, M. Caparon, and S. J. Hultgren (2015). Urinary tract infections: epidemiology, mechanisms of infection and treatment options. Nature reviews microbiology 13.5, pp. 269– 284.

33. A. Vasala, V. P. Hytönen, and O. H. Laitinen (2020). Modern tools for rapid diagnostics of antimicrobial resistance. Frontiers in Cellular and Infection Microbiology 10, p. 308.

34. E. J. Gauger, M. P. Leatham, R. Mercado-Lubo, D. C. Laux, T. Conway, and P. S. Cohen (2007). Role of motility and the flhDC operon in Escherichia coli MG1655 colonization of the mouse intestine. Infection and immunity 75.7, pp. 3315– 3324.

35. F. Wu, A. Japaridze, X. Zheng, J. Wiktor, J. W. Kerssemakers, and C. Dekker (2019). Direct imaging of the circular chromosome in a live bacterium. Nature communications 10.1, pp. 1–9.

36. A. Japaridze, C. Gogou, J. W. Kerssemakers, H. M. Nguyen, and C. Dekker (2020). Direct observation of independently moving replisomes in Escherichia coli. Nature communications 11.1, pp. 1–10.

37. A. Castellanos-Gomez, R. van Leeuwen, M. Buscema, H. S. van der Zant, G. A. Steele, and W. J. Venstra (2013). Single-layer MoS2 mechanical resonators. Advanced Materials 25.46, pp. 6719– 6723.

38. B. Volkmer and M. Heinemann (2011). Condition-dependent cell volume and concentration of Escherichia coli to facilitate data conversion for systems biology modeling. PloS one 6.7, e23126.

39. M. Riley (1999). Correlates of smallest sizes for microorganisms. In Size limits of very small microorganisms: proceedings of a workshop. Vol. 3. National Academies Press Washington DC, USA, p. 21.

