## Supplementary Notes 1-6 for "Microwell-enhanced optical rapid antibiotic susceptibility testing of single bacteria"

### CONTENTS

Supplementary Note 1: Extended measurement data 2
Supplementary Note 2: Confirmation of weak trapping in wells by microscopy using transparent PDMS patterns. 3
Supplementary Note 3: Extended simulation data from COMSOL of a laser beam tightly focused on a bacterium. 4
Supplementary Note 4: Influence of antibiotics on cell morphology 5
Supplementary Note 5: Detecting antibiotic resistance with Micro-wells 6
Supplementary Note 6: Role of motility in signal detection 7
Supplementary Note 7: Measurements on 2.5 micron deep micro-wells 8
References 8

*

### SUPPLEMENTARY NOTE 1: EXTENDED MEASUREMENT DATA

In the main text the results of measurements on bare silicon and in etched wells are described. Here, we show typical traces for each of the cases as a supplement to the data presented in the main text.

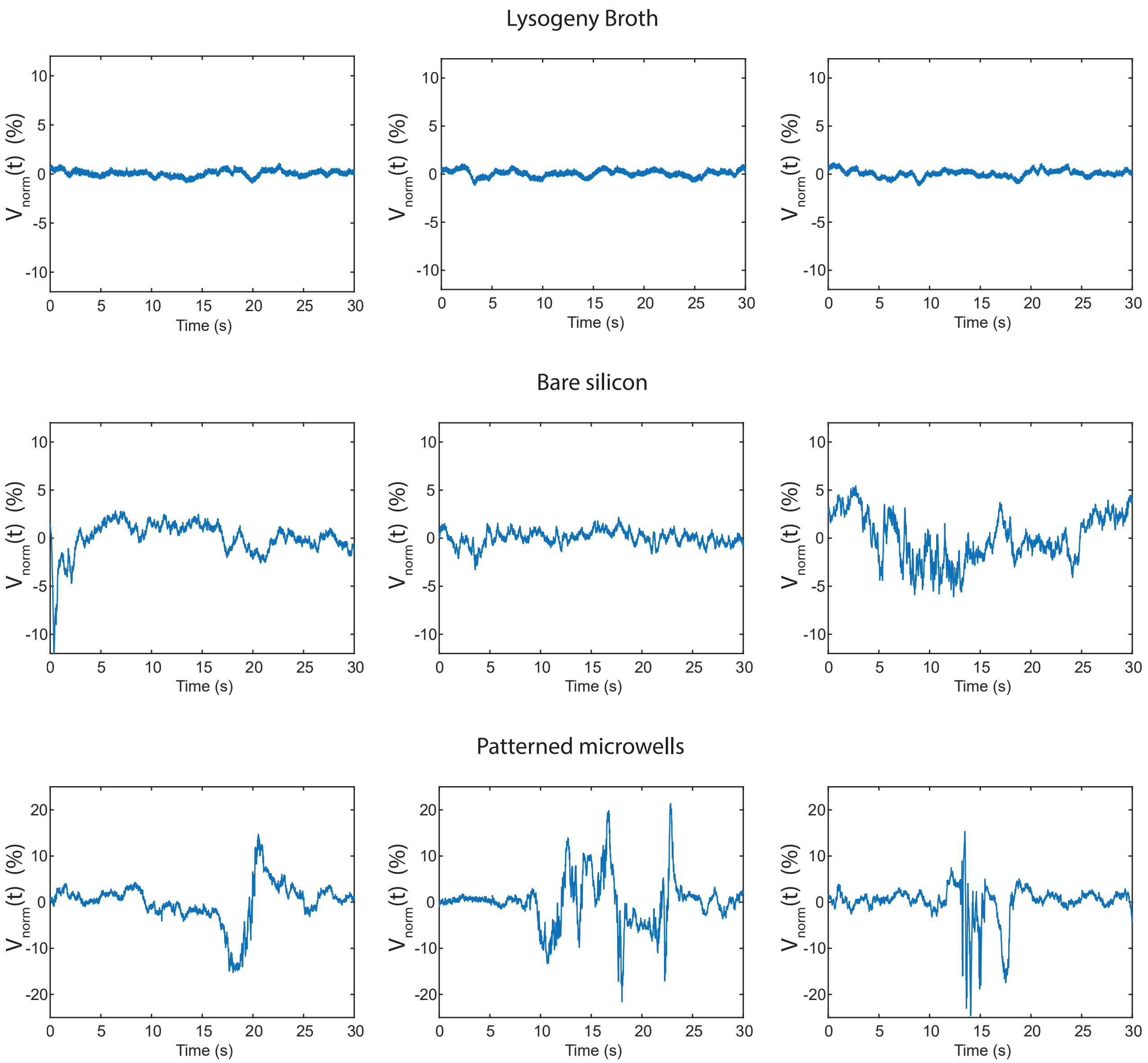


FIG. S1. Impact of substrate on observed signal, extended graphs to figure 2 in the main text. Each graph shows a measurements performed on single position or microwell during 30 seconds.

### SUPPLEMENTARY NOTE 2: CONFIRMATION OF WEAK TRAPPING IN WELLS BY MICROSCOPY USING TRANSPARENT PDMS PATTERNS.

PDMS substrates, with the same geometry as the silicon chips used in the main text, were prepared to validate microscopically that the cells are weakly trapped within the 285nm deep wells by measuring residence times inside the wells with respect to outside. The moulded PDMS cavities are transparent, allowing for the use of a transilumination microscope to visualize the cells. These samples were placed under a Nikon Ti-E microscope with a 100X CFI Plan Apo Lambda Oil objective with an NA of 1.45 equipped with a phase ring. Videos of 120s were recorded to observe the residence of the cells. These videos were analyzed using a MATLAB script to track the cells and compare occupation of areas inside and outside of the wells. A raw video frame and the result of the cell recognition algorithm are shown in figure S2a. The areas where cells are present are marked white. Then, a mask is applied to find if a cell is inside or outside of a well. For each movie frame, we calculated the cell count normalized by the mask area, $A_{\rm cell}/A_{\rm mask}$, both inside and outside of the wells, as shown in figure S2b and c. The occupation of wells is approximately 50% higher than of the surrounding area, confirming a weak trapping of the cells.

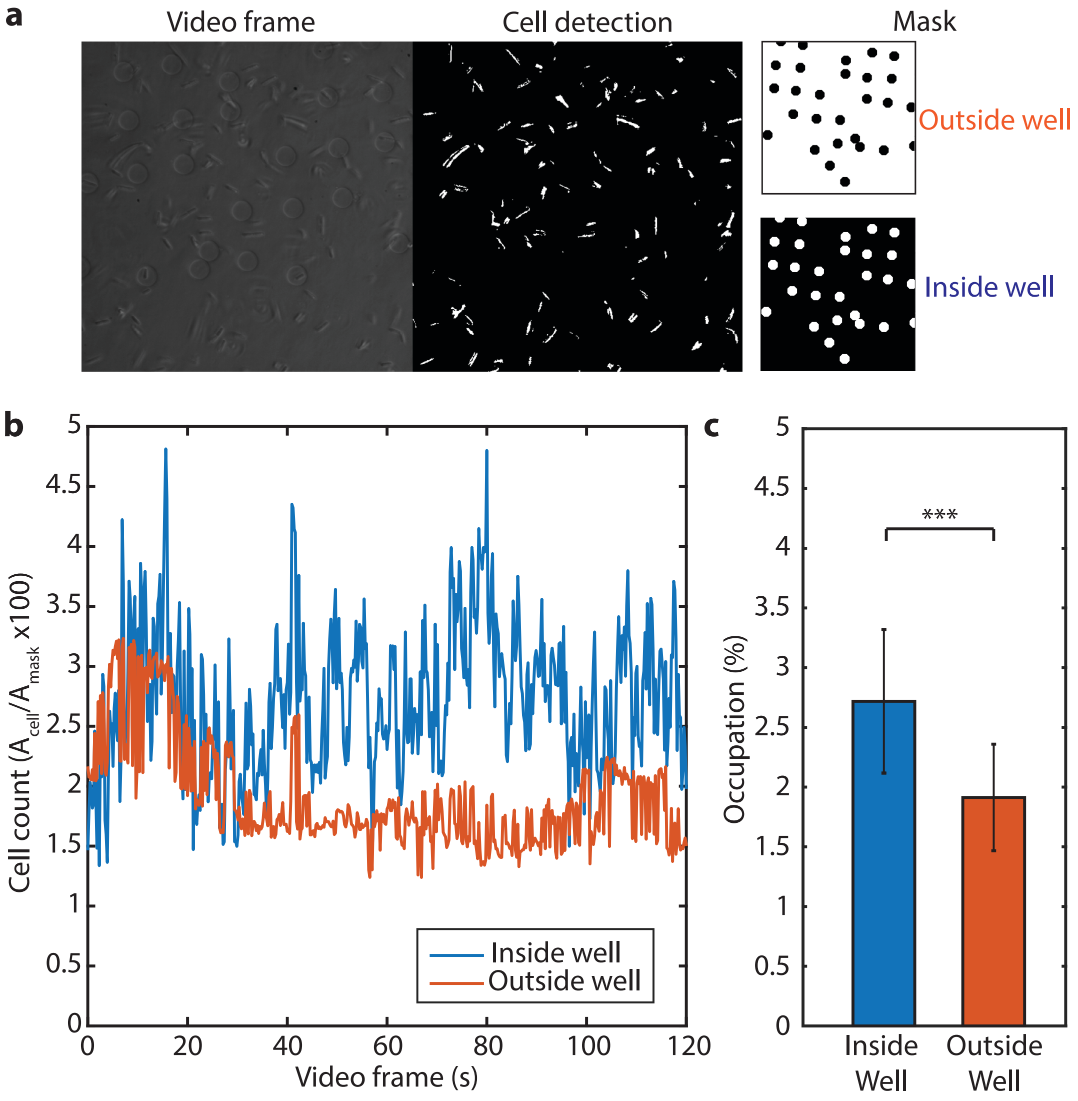


FIG. S2. Occupation of 285nm deep PDMS wells by *E.coli* cells. a) A raw video frame and the result of the cell detection algorithm used for analysis. b) The percentage of the total area that is occupied by cells is calculated both inside and outside of wells for the entire length of the video. c) The occupation of wells is approximately 50% higher than of the surrounding area.)

### SUPPLEMENTARY NOTE 3: EXTENDED SIMULATION DATA FROM COMSOL OF A LASER BEAM TIGHTLY FOCUSED ON A BACTERIUM.

COMSOL simulations were performed to estimate the light scattered and absorbed by a single bacterium in a tightly focused beam. Here, we show several beam profiles at varying lateral position of the bacterium, which were used to obtain figure 3c and 3d in the main text.

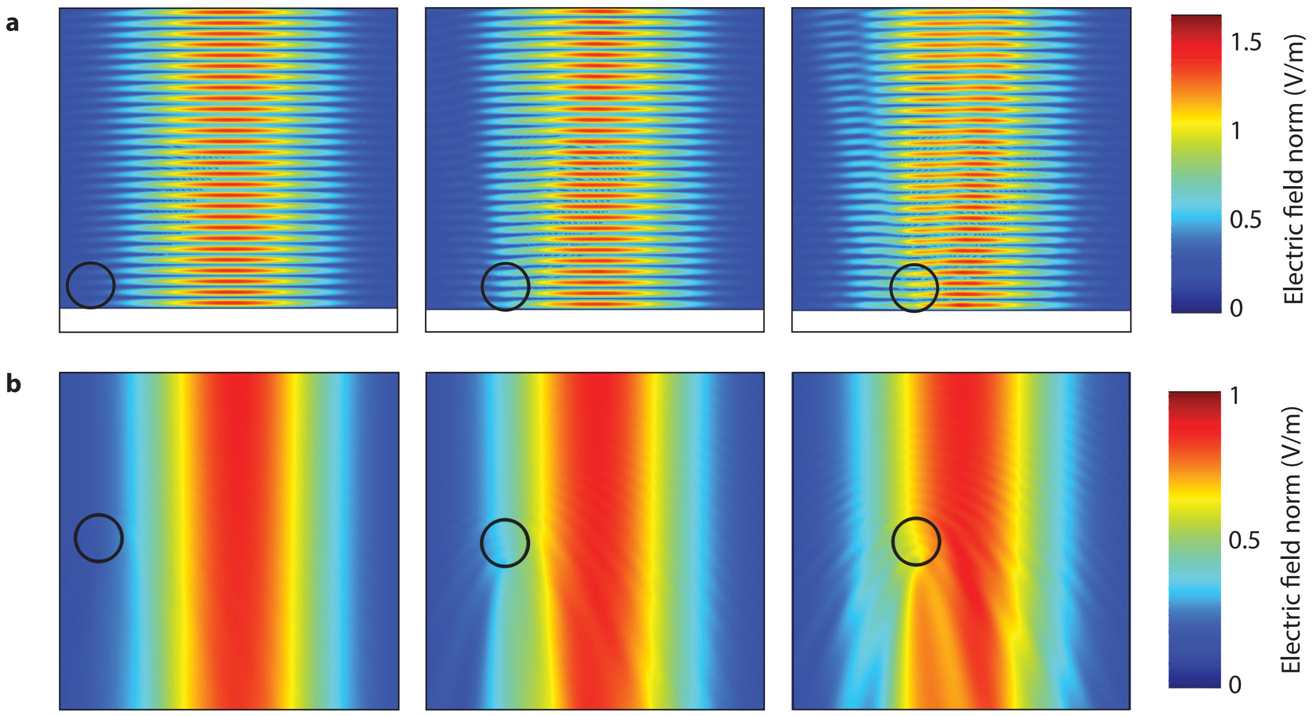


FIG. S3. Simulation of a circular bacterium at varying lateral position with respect to a tightly focused laser beam. Extended simulation results to figure 2 in the main text are shown of the electric field norm $E$, at lateral positions $x = -3$,$x = -2$, and $x = -1\mu$m with respect to the centre of the beam.

### SUPPLEMENTARY NOTE 4: INFLUENCE OF ANTIBIOTICS ON CELL MORPHOLOGY

Experiments were performed on *E.coli* cells in the presence of ciprofloxacin and chloramphenicol antibiotics. Bacteria were grown under agarose pads to ensure they stay within the imaging field of view as previously described by A. Japaridze et al 2020. The addition of antibiotics did not influence the rodshape of the bacteria and its phenotype.

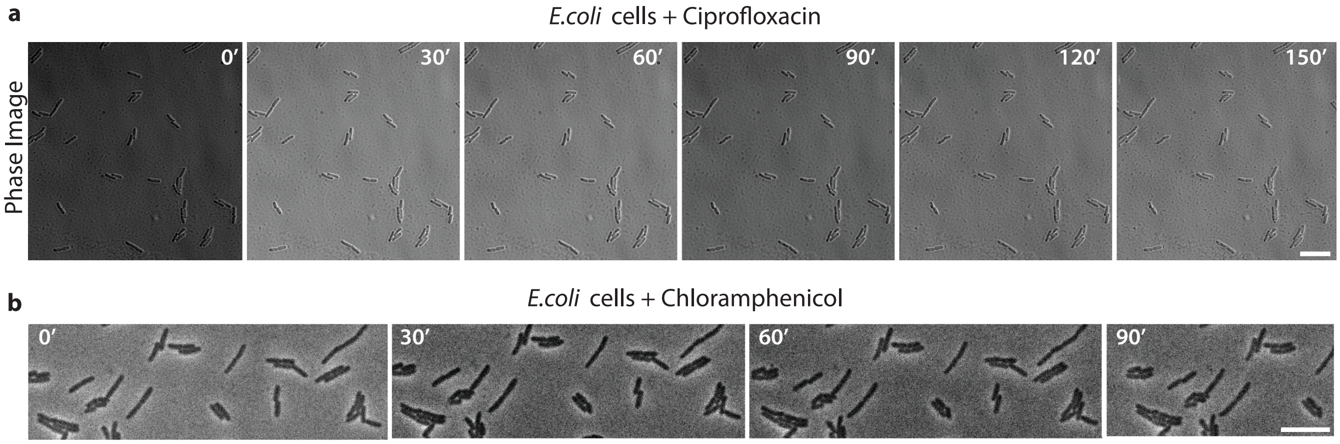


FIG. S4. Phase contrast images of cells grown on LB under agarose pads. After exposure to either the Ciprofloxacin a. or Chloramphenicol drug, b. the morphology of cells does not change. Time indicated in minutes. Scale bars, 10 microns.

### SUPPLEMENTARY NOTE 5: DETECTING ANTIBIOTIC RESISTANCE WITH MICRO-WELLS

Experiments were performed on *E.coli* cells with a chromosomal *KanR* resistance gene exposed to Kanamycin. As it can be observed in figure S6, no change in the signal variance is seen after exposure to antibiotic, showing that the platform can potentially be used for fast detection of antibiotic resistance with single cell resolution.

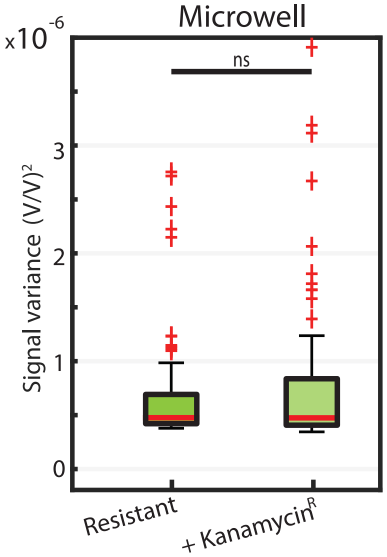


FIG. S5. Signals before and 1 hour after administering kanamycin (25$\mu$g/ml final concentration) to MG1655(*kanR*) resistant cells. There is no signal drop observed for kanamycin resistant strains ((light green, ($n = 84$)($p = 0.53$, ns))

### SUPPLEMENTARY NOTE 6: ROLE OF MOTILITY IN SIGNAL DETECTION

Experiments were performed on *E.coli* cells lacking the motA and motB genes resulting in non-motile cells. When the signal for the non=motile bacteria was measured, it was comparable to the background signal. Even addition of meropenem antibiotic (50$\mu$g/ml 1.5 hour after exposure) that the cells were susceptible to, did not reduce the signal variance, indicating that the the signal drop was only observed for motile cells. We used a rank sum test for comparison between the conditions, with the following convention ns: $0.05 < \text{p}$.

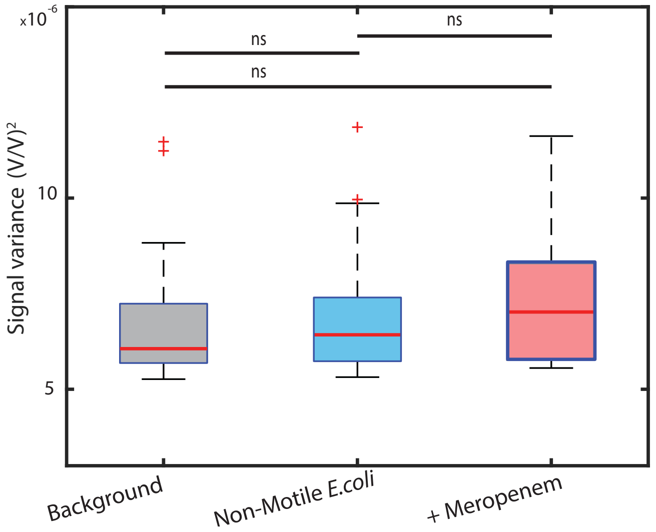


FIG. S6. Signals on bare chips, before and 1.5 hour after administering meropenem (50$\mu$g/ml final concentration) to *E.coli motAB* non-motile cells. There is no statistical difference between the signal on empty wells ((light grey, ($n = 41$), as well as on non-motile ((light blue, ($n = 71$) or antibiotic treated cells ((light red, ($n = 82$).

### SUPPLEMENTARY NOTE 7: MEASUREMENTS ON 2.5 MICRON DEEP MICRO-WELLS

We also performed experiments on deeper micro-wells to verify whether the depth might have influence on trapping. For this purpose we prepared micro-wells that are 2.5 micron deep, rather than 285 nm that was used in the main text. As it can be observed in figure S7, trapping events that last more than 10 seconds occur in the deeper wells, during which signal fluctuations can be recorded.

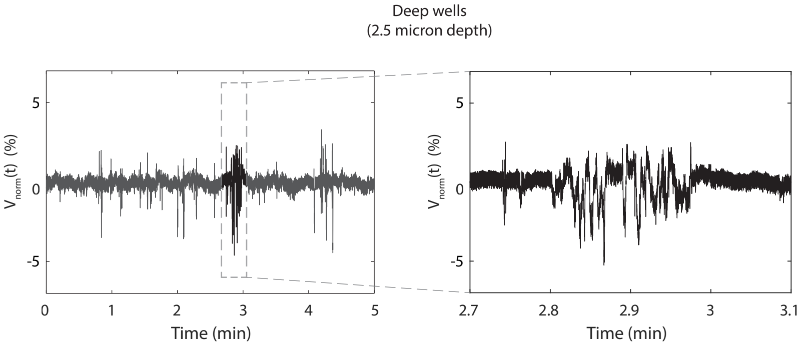


FIG. S7. Signals recorded 2.5 micron deep microwells. The signal was recorded for 5 minutes. A trapping event is indicated on the left panel, and the signal is shown in higher detail on the right.
